# Identifying planktonic foraminiferal morphospecies: which are the important images?

**DOI:** 10.1101/2021.04.13.439602

**Authors:** George H. Scott

## Abstract

Selection of imagery that promotes accurate identification of morphotaxa is viewed as a significant problem in the taxonomy of planktonic foraminifera. Currently, imagery of taxa is sparse, apparently selected by visual judgement, and presented without information about its typicality. What is required are impartially selected images which embrace population variation to serve as training sets for reliable identification of taxa. Outlined here is a simple morphometrically-based solution, applied to the shape of shells in two orientations, in which shape variation is resolved onto three principal component axes. On the premise that the best-adapted shells are the commonest, specimens within 1 standard deviation (sd) of the trivariate mean are recognized as population exemplars suitable for use as trainers. Specimens which project at ≥2 sd onto at least one axis are mapped as boundary specimens whose identity might be questioned. This procedure is trialled on samples of *Truncorotalia crassaformis*. Exemplars from the Equatorial Atlantic and Caribbean compare closely; they partially overlap with those from a Holocene Southwest Pacific population provisionally interpreted as a subtaxon, *Truncorotalia crassaformis hessi*.

## Introduction

Identification implies a taxonomy and for experts and tyros alike it is often necessary to refresh the visual concept of taxa. For this purpose a default reference for foraminifera has long been Ellis & Messina (2008) which is a compilation of taxon proposals extracted from the literature. A sample of the literature (Scott, 2011) shows that few specimens other than the holotype are illustrated. This specimen is a name-bearer and is obligatory in zoological taxonomy to stabilize nomenclature, although the Code of Zoological Nomenclature does not demand its illustration. Historically, a focus on the holotype as the sole exemplar of a species may reflect a typological approach to taxonomy or the constraints of manual artistry. Regardless, modern optical and electron microscopy obviate any need for a minimalist approach to species imagery. Yet, in this age where digital sources are fast replacing print, imagery of planktonic taxa in online databases and literature remains small. For *Truncorotalia crassaformis*, studied here, Schiedel and Hemleben (2017, pl. 2.23) show three views of two specimens and four single images; WoRMS (http://www.marinespecies.org/) shows two specimens; microtax (http://www.mikrotax.org/pforams/) shows 10 specimens in three orientations supplemented by five single images; Foraminifera.eu (http://www.foraminifera.eu/) shows two views of 4 specimens and 2 single views. Outstanding is http://endless.forams.org which shows one view of 287 specimens.

While the power of imagery for taxonomy is under-utilized, what is the status of these images? They might be recognized as plesiotypes: specimens, other than the type, upon which a subsequent or supplementary description or figure is based (syn: apotype, heautotype, plesiotype; Evenhuis, 2008). However, these are informal categories not recognized in the Code of Zoological Nomenclature. Here they are called exemplars: specimens that authors regard, almost always implicitly, as representative of a species. As such, their imagery is important for the transfer of information about a species as well as allowing others to assess an author’s interpretation of it. In this way it conforms with a tenet of scientific research and the principal complaint might be only that there is insufficient such imagery (Boltovskoy 1965; Hsiang et al., 2019). Where it is open to further examination is the selection of imagery. How did the author decide that the specimens are representative? This is seldom addressed and is likely to have been made by visual judgement. Is that procedure fit for purpose and is it adequate for accurate identification of the species? In practice, which imagery should be presented to viewers that best facilitates specimen identification?

Molecular studies (Darling & Wade, 2008; Quillévéré et al., 2013; Andre et al., 2014) show that the distribution of populations is watermass related, suggesting that the search for morphogroup exemplars should be similarly based on sampling of local populations, site by site. Which specimens should be selected? As an approximation, it is proposed that the best adapted morphotypes are the commonest in living and, more equivocally, in populations preserved in bottom sediments. This suggests that specimens near the center of a sampled population should the most informative architectural exemplars (C-exemplars). At the cognitive level this aligns with experimentation (Iordan et al., 2016) which shows that the neurovisual system performs best with typical exemplars. Viewed simply as objects to be identified, what features are the most informative? There is consensus in the neuroscience community that shape is an especially powerful attribute (Elder, 2018 and references therein). As much information on the shape of an object is encapsulated in its 2D outline, the distribution of this attribute is used here as data for identifying C-exemplars. But what of the atypical specimens? These present an important problem for those enumerating taxa for their use as environmental proxies (Schiedel et al., 2018). A partial solution is to identify specimens at the boundary of a site distribution mapped for C-exemplars as B-specimens. These procedures are applied to modern and Holocene populations of *Truncorotalia crassaformis*, a species that illustrates the variability in shell shape commonly found in planktonic foraminifera (Lidz, 1972; Lamb and Beard, 1972; Bolli and Saunders 1985).

## Material and Methods

DSDP Site 366A 1-1W-3-5 cm. (ETA): This site (05° 40.7 N; 19° 51 W; 2853 m) is on the Sierra Leone Rise in the eastern tropical Atlantic and lies under the Equatorial Counter Current. It is near Core 234 examined by Lidz (1972). From the model of Lazarus et al. (1995), the age of the sampled horizon is <3 kyr. Thirty-five specimens from the > 149 μm fraction were extracted by the author.

Gulf of Mexico sediment trap (GM): This trap is a time series (2008–2012) of foraminiferal and particulate flux at 700 m on the northern Gulf of Mexico continental shelf (27.5° N; 90.3° W). Dr. Caitlin Reynolds supplied 50 specimens (212 μm–425 μm fraction) from the GMT2-1 sediment trap (Richey et al., 2014). Specimens were collected between 21–27 April, 2008.

DSDP Site 591A 1-1-1 cm (SWP): This site (31°35.06 S; 164°26.92 E; 2142 m) is on a southern spur of the eastern part of the Lord Howe Rise in the vicinity of the Tasman Front, a westward flowing branch of the East Australian Current. The age of the sample (Nelson et al.,1993) is 9.4 kyr; 30 specimens were extracted from the >149 μm fraction by the author.

### Methods

Sediment residues from ETA and SWP were strewn on a gridded tray and globorotaliid specimens with <5 chambers in the outer whorl and a narrow aperture were selected as encountered. These criteria exclude taxa such as *Truncorotalia truncatulinoides, Globorotalia menardii* and *Globoconella inflata* but admit specimens that closely resemble *Truncorotalia crassaformis*, e.g., *Truncorotalia crassula*. Dr. Catlin Reynolds selected GM material. Specimens from the three localities were imaged under scanning electron microscopy in standard axial and spiral orientations. Outline data were manually captured (tpsdig2, http://life.bio.sunysb.edu/morph/soft-dataacq.html) as 180 equally-spaced coordinates. Raw data were processed using generalized procrustes analysis (GPA, http://cran.r-project.org/web/packages/shapes/index.html) which aligns specimens on their centroids and removes size and positional differences (Webster and Sheets, 2010). Principal component (PC) projection of the high-dimensional data onto 3 orthogonal axes retains 63-82% of the shape information and allows inspection of the specimen configuration in low dimensional euclidean space. As the sampling model potentially allows for multiple groups their presence is checked using fuzzy clustering (Ferraro et al. (2019). In samples interpreted as representing a single morphogroup C-exemplars are those within 1 standard deviation (1 sd) of the trivariate mean of the standardized PC scores. B-specimens are those individuals in which one of the three PC scores is greater than 2 sd of its mean.

## Results

‘Normalform’ and ‘n-form’ refer to specimens in which, in late ontogeny, chambers increase in size with little change in shape. ‘Kummerform’ and ‘k-form’ refer to specimens in which the last-formed (fth) chamber is dimensionally smaller than its predecessor (f-1th).

Recall that specimens are selected only from data on their outline shape; other data in the images are ignored. The number of C-exemplars and B-specimens per sample reflects the distribution of its PC scores and the grouping levels. Following experimental neuroscience research (Alvarez, 2011; Whitney & Leib, 2018), C-exemplars are shown as sets of close-packed images at a standard size to enhance their value for visual recall, comparison with published images and with observed specimens.

### Axial Shape

The trivariate plots (Figs 2–4) show that the B-specimen criterion (1 PC variable with standardized score >2 sd from its mean) identifies marginal specimens that vary in shape from elliptical (Fig. 2 #a16) to cone-like (Fig. 2 #a23) thus spanning shapes seen in the more tightly constrained C-exemplars. Elliptical shapes (strong extension of the outline radially) develop in specimens in which the last-formed (fth) chamber is smaller than its predecessor (f-1 th; kummerform condition). More compact, cone-like shapes arise when the fth chamber is larger than the f-1th and extends strongly in the direction of the coiling axis. Axial extension in some is greater than in the associated C-exemplars. Lesser contributors to shape variation in C-exemplar and B-specimens are the height of the spire formed by early whorls, curvature of the umbilical wall of the fth chamber, the angularity of the junction with its spiral wall and the orientation of the latter. Some extreme shapes (Fig. 4 #a21) are created by the size and orientation of a kummerform fth chamber and are interpreted as intra-population variants, some other B-specimens which map as shape outliers (Fig. 4 #a25) have grown normally and their relation to the population might be questioned.

**Fig. 1.**
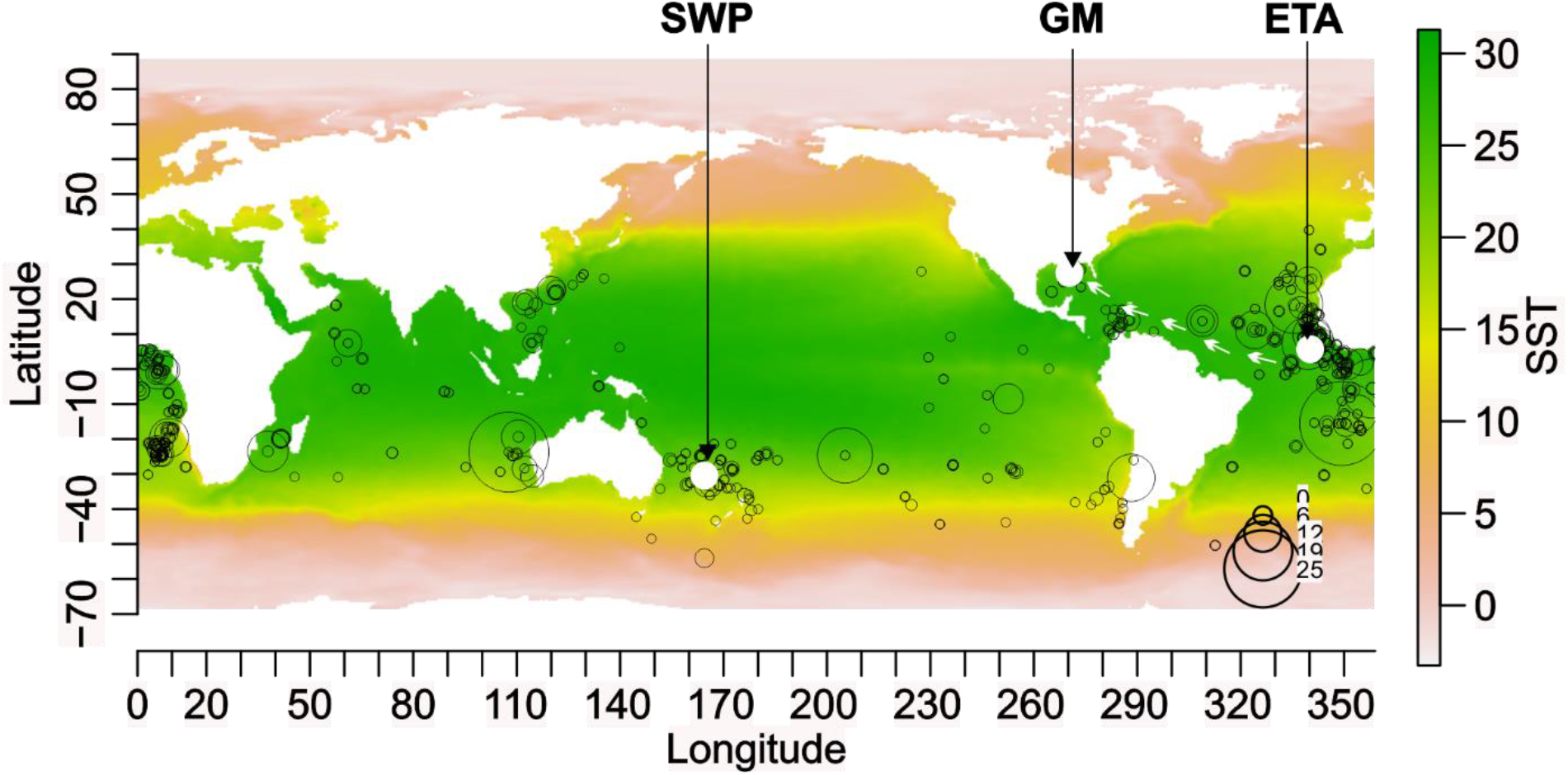
Distribution of *Truncorotalia crassaformis* in the coretops database ForCenS (Siccha & Kučera, 2017) plotted on global sea surface temperatures from CARS 2009 (Ridgway et al., 2002). Records in which the species abundance ≥ 2% of an assemblage are plotted.

**Fig. 2.**
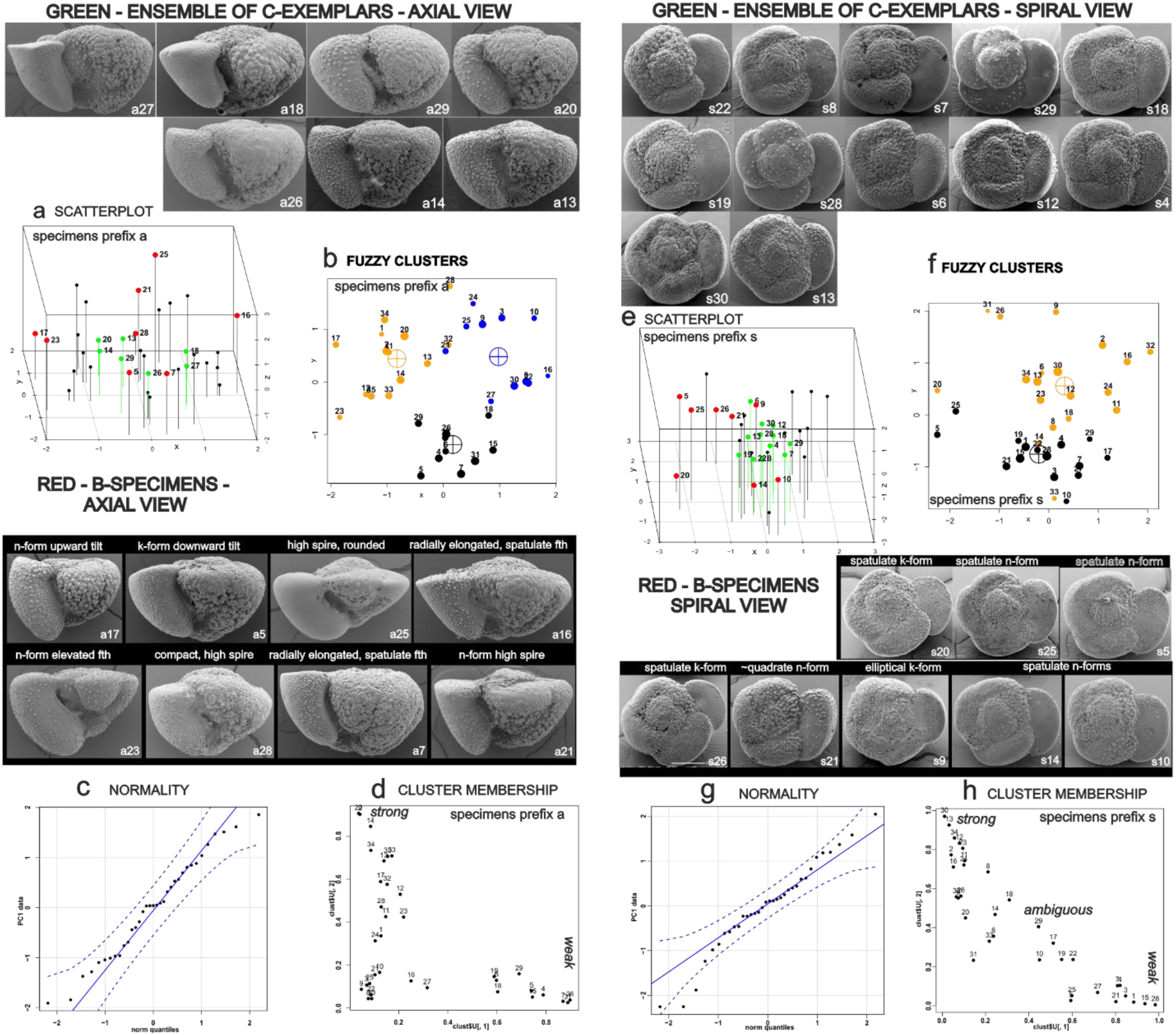
Variation in shape using outline data for *Truncorotalia crassaformis* in axial and spiral orientations from DSDP Site 366A 1-1W-3-5 in the equatorial Atlantic (ETA in the text). (a, e) Positions of C-exemplars and B-specimens from PCA projections of the axial/spiral outline data. (b, f) 3D scatterplots identifying C-exemplars (green) and B-specimens (red) (c, g) PC1 QQ normality, envelope = 0.95 confidence level. (d, h) Degree of cluster membership: points ~(1,0) = strong membership; ~(0.5,0.5) ambiguous membership; ~(0,0) = low membership. The R-specimen criterion (red in (a, e) detects most of the outliers in either orientation. More specimens identify as C-exemplars in the spiral data (e) but cluster analysis partitions them between two groups, discriminating spatulate specimens (e.g., #a7, #a22) from normalforms (e.g., #a6, #a12). Differences in the shape of late-formed chambers account for the poorly defined clusters in both axial and spiral data. The sample is interpreted as representing a single morphogroup which exhibits large shape differences in late ontogeny. Specimen #a25 possibly represents *Truncorotalia crassula*.

**Fig. 3.**
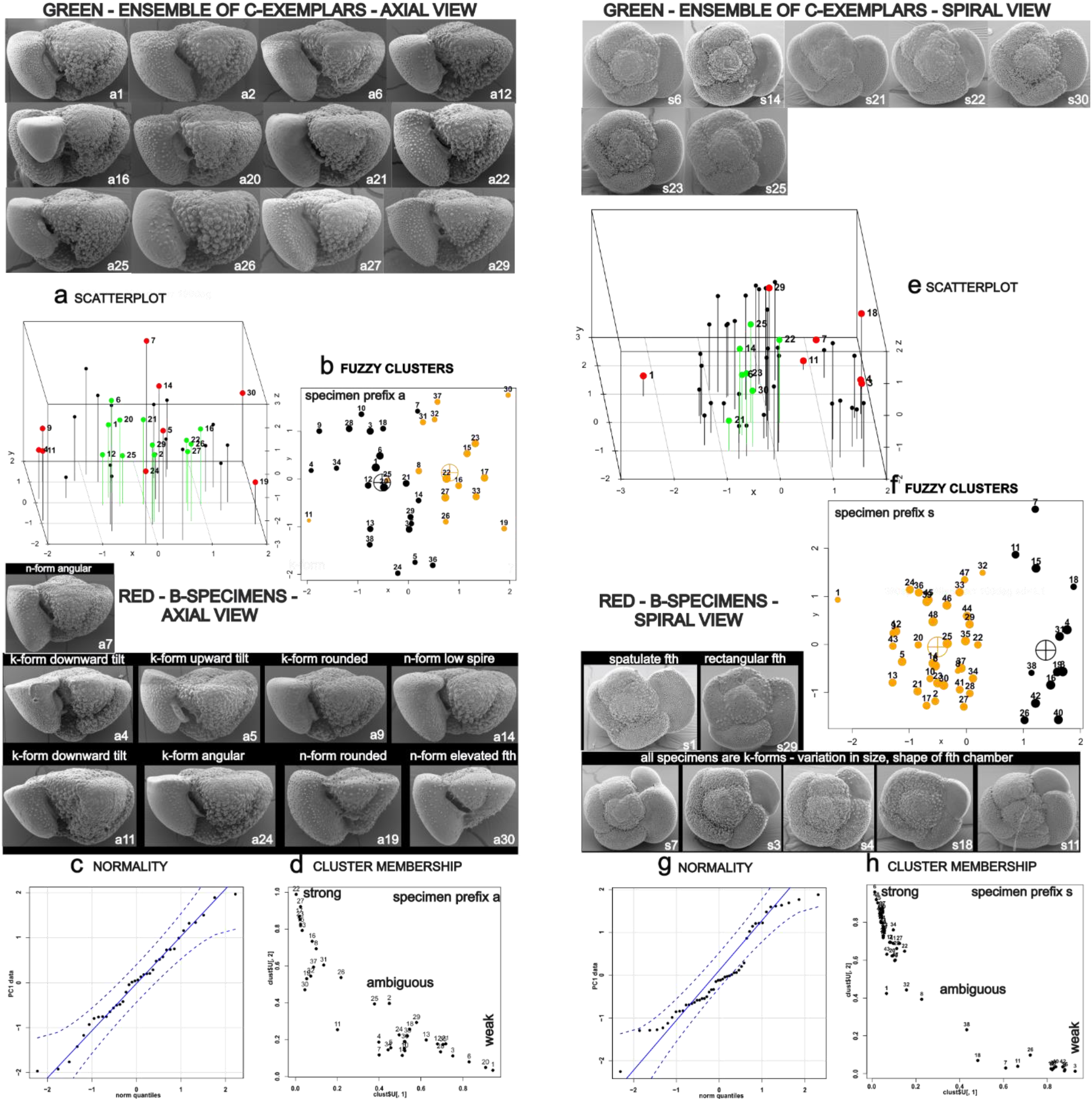
Variation in shape using outline data for *Truncorotalia crassaformis* in axial and spiral orientations from the Gulf of Mexico sediment trap (GM in the text). (a, e) Positions of C-exemplars and B-specimens from PCA projections of the axial/spiral outline data. (b, f) 3D scatterplots identifying C-exemplars (green) and B-specimens (red) (c, g) PC1 QQ normality, envelope = 0.95 confidence level. (d, h) Degree of cluster membership: points ~(1,0) = strong membership; ~(0.5,0.5) ambiguous membership; ~(0,0) = low membership. C-exemplars (a) are widely dispersed in the axial data and the clusters (b) are poorly discriminated with many specimens whose classification is ambiguous. Membership is better defined in the spiral data and C-exemplars are in closer proximity: this reflects the lesser presence of spatulate specimens than in Fig. 3. They are recognized here as R-specimens (outliers). The low level of cluster membership in the axial data supports acceptance of the sample as one from a single morphogroup.

**Fig. 4.**
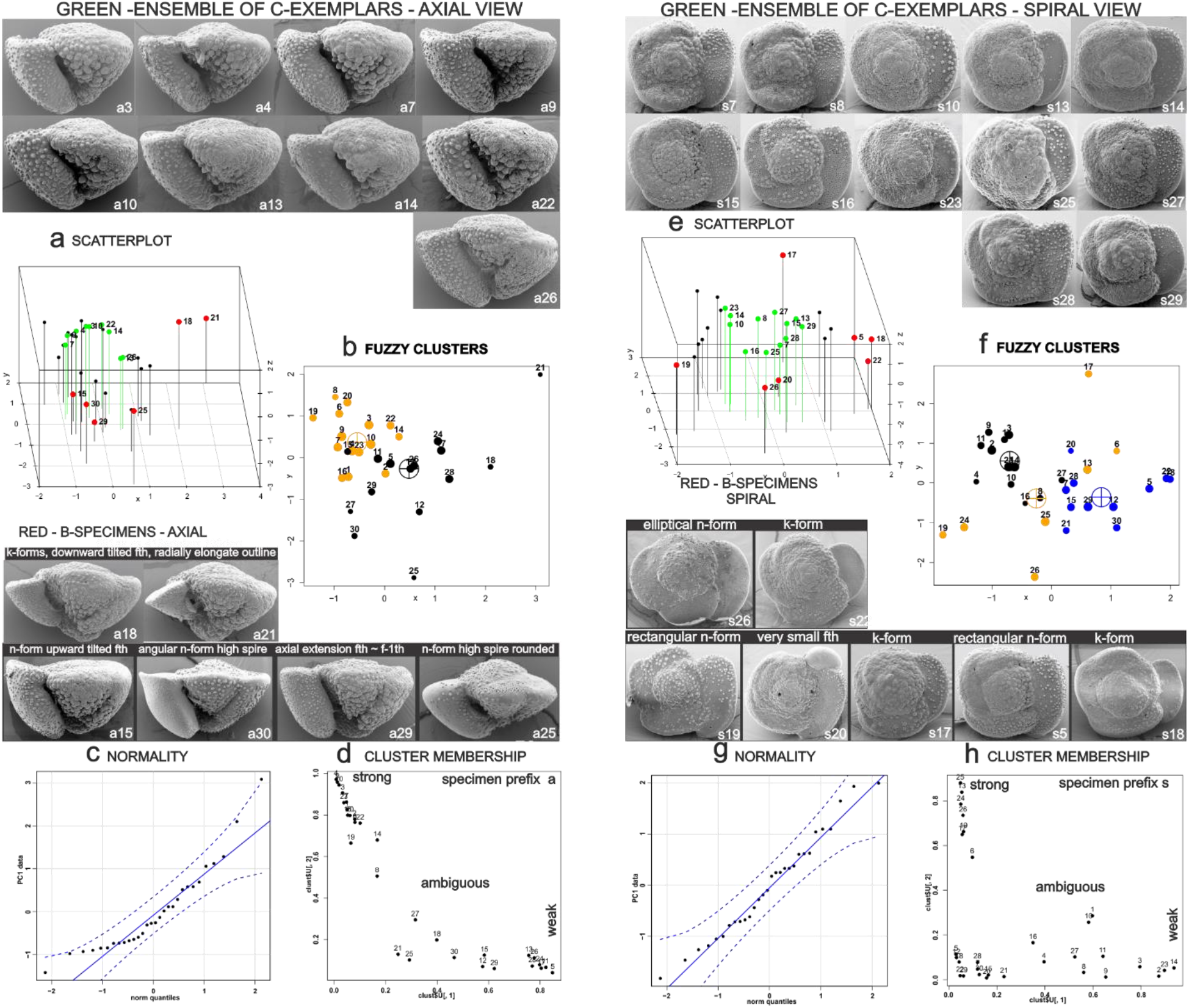
Variation in shape using outline data for *Truncorotalia crassaformis* in axial and spiral orientations from DSDP Site 591A 1-1-1 in the Southwest Pacific (SWP in the text). (a, e) Positions of C-exemplars and B-specimens from PCA projections of the axial/spiral outline data. (b, f) 3D scatterplots identifying C-exemplars (green) and B-specimens (red). (c, g) PC1 QQ normality, envelope = 0.95 confidence level. (d, h) Degree of cluster membership: points ~(1,0) = strong membership; ~(0.5,0.5) ambiguous membership; ~(0,0) = low membership. Visually, axial C-exemplars are closely similar in shape: all are normalforms. B-specimens include #a15 a normalform in which the spiral surface of the fth chamber is angled upward and rises above the early whorls; similar architecture is found in *Truncorotalia truncatulinoides*. C-exemplars (spiral) include good examples (#s8, #s28) of the distinctive elongated, low-curvature outline of the fth chamber. In both axial and spiral data many specimens have weak or ambiguous cluster membership. The sample is considered to represent a single morphogroup. A caveat is specimen #a25, a normalform outlier whose shape is similar to that of #a25 in Fig. 3.

### Spiral Shape

This view (Figs 2–4) is a slice through a trochoidally-coiled shell at the spiral surface of the outer whorl. Its outline, which varies from nearly circular through elliptical to subquadrate, is modulated by the size and shape of all chambers in that whorl. Brief inspection of C-exemplars (Fig. 2–4) suggests little difference in inter-sample shape. Closer examination shows that subquadrate outlines are best developed in some SWP (Fig. 4) exemplars while elliptical outlines are best developed in some ETA (Fig. 2) exemplars. The shape and especially the outline of late-formed chambers contributes to this difference. In some SWP exemplars these chambers extend in the direction of growth with low curvature along their outer margin. Figure 4 #s19 is a extreme example. In contrast, in some ETA and GM C-examplars and B-specimens curvature along the margin of the fth chamber is much greater than in its predecessors producing spatula-like shapes. It is best-developed in kummerforms (Fig. 2 #s20, #s26; Fig. 3 #s1, #s3). Nevertheless, there are rare specimens in ETA and GM in which the fth chamber resembles the low curvature outline common among SWP C-exemplars.

### Discrimination

As well as partitioning the shape data for visual objectives it is relevant to map them as multivariate populations. The canonical discrimination plot for axial outline data (Fig. 5) shows that GM and ETA specimens closely overlap and that their means overlap at the 95% confidence level. The scatter of SWP specimens partially overlaps those for GM and ETA but its mean is clearly distinct. The greater presence of kummerform specimens in the GM sample increases its separation from the ETA scatter for the spiral outline data.

**Fig. 5.**
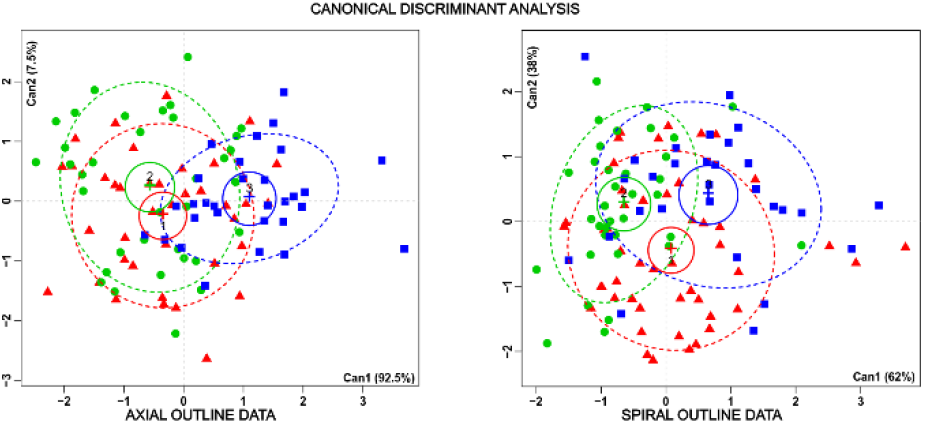
Generalized canonical discriminant analysis (candisc package, available from https://cran.r-project.org/web/packages/)using a multivariate linear model on PC1-3 axial and spiral data for Gulf of Mexico sediment trap specimens (red, GM in text), DSDP Site 366A 1-1W-3-5 in the equatorial Atlantic (green, ETA in the text) and DSDP Site 591A 1-1-1 in the Southwest Pacific (blue, SWP in the text). Confidence intervals for means (95%) and data ellipses (68%) are shown.

## Discussion

### Caribbean (GM) and tropical Atlantic (ETA) populations

Although compromised by differences in sampling (coretop:sediment trap) and selection of specimens, comparison of data on shell shape from these localities is relevant because surface currents (Fig. 1 white arrows) suggests that they might sample a common biological population. On that assumption the comparison examines the stability of outline shape in a water mass over a large distance. Visually, their axial C-exemplars are comparable; this is supported by the multivariate analysis (Fig. 5) of the full sample data which shows that their confidence intervals for mean shape overlap, as do their canonical scores. The close similarly in axial shape, both visually and statistically is weakened in the spiral data, principally by the greater presence of specimens with spatula-like fth chambers in the ETA population. Again, the statistical difference is supported by the C-examplars for ETA which include such specimens. Possibly the difference may be related to the greater depth of the ETA sample which, as a bottom sample, covers the full depth range of populations in the upper ocean.

### Southwest Pacific (SWP) population

This population might perplex identifiers: it includes some specimens similar to those in ETA and GM but has others that are distinctive. This is a common problem in the classification of planktonic foraminiferal morphospecies that has long been recognized in *Truncorotalia crassaformis* (Parker, 1962; Lidz, 1972; Gradstein, 1975; Stainforth et al., 1975; Bolli and Saunders, 1986; Bylinskaya 2005). Visually, C-exemplars for the axial outline in SWP are the most distinctive set in this study. This is supported by statistics for the full sample which show that mean axial shape is distant from the GM and ETA populations. While wide shape variation among the axial B-specimens might puzzle identifiers, the C-exemplars support recognition of the SWP population as a morphogroup separable from that in GM and ETA. Northward from SWP Chaproniere (1991) identified similar specimens as *Globorotalia crassaformis hessi*, proposed as *Globorotalia hessi* by Bolli and Premoli Silva (1973) from Caribbean DSDP Site 19. Several images (e.g., ZF5888-Globorotalia-crassaformis_obj00008_plane000.jpg in http://endlessforams.org/) have the quadrate outline and subrectangular chambers seen in some SWP specimens and also come from a site in the South Pacific Subtropical Gyre. Whether the *hessi* morphogroup is present in the modern gyre is unresolved but its unwitting inclusion with *Truncorotalia crassaformis* in Quaternary censii might affect the use of the latter as an environmental proxy.

### Samples and Training Sets

In the language of artificial intelligence (AI) C-exemplars are training sets for the visual identification of taxa. Whether directed to automatic or visual identification the composition of the training set should mirror attributes of the morphogroup it samples. This raises questions: how to define the taxon as a morphogroup and how to sample it. In the absence of genomic data the approach here is to define characters shared by a wider morphogroup, apply those criteria to a strew of an assemblage (living/dead) and examine the potentially multi-taxon sample for its integrity as a single morphogroup, assisted by cluster analysis of the shape data. This enhances the transparency of the selection of specimens for possible use in a training set although expert opinion is involved in interpretation of the cluster analysis. Given the morphogroup this study recognizes specimens within 1 sd of the trivariate mean as exemplars. This is a form of purposive sampling (Etikan et al., 2016) used here to locate typical specimens: those that, following https://www.lexico.com/definition/Typical, have the distinctive shape attributes of the morphogroup. Use of this algorithm removes operator bias but assumes that sampling is from a normal distribution. This applies to most of the data in this study (Fig. 2–4) but note that definition of the C-exemplar set requires operator choice of the sd value. There are several strategies for detection of R-specimens (e.g., Routliers, available at https://cran.r-project.org/web/packages/Routliers/index.html) that might be used.

Hsiang et al. (2019) take a quite different approach to the recognition of exemplars wherein a panel of 24 expert taxonomists identified taxa from images of specimens selected from 35 Atlantic coretops. Those images for which there was 75% agreement amongst at least 4 experts were retained as training sets. Their procedure represents consensus judgement by experts based on pooled samples whereas that trialled here uses site-specific sampling and algorithmic identification of exemplars. Both strategies attempt to reduce selection bias but the algorithmic approach applied here offers greater transparency.

### Implementations

A theme of this study is the need to sample and illustrate populations in ways that represent population variation, raise transparency in specimen selection and reduce operator judgements. Identification errors reported by Fenton et al. (2018) underline the need for research on the theme. As the procedure trialled here might be regarded as unsuitable because of its use of morphometric data and algorithms for selection, simpler methods might be considered. For example: split a residue down to a volume that fits the viewing tray and sample *Truncorotalia crassaformis* in a way which allows every specimen an equal chance of selection. Exemplars would span the shape population with common architectures well represented and outliers seldom represented. Apart from the initial decision by the operator about the identity of specimens, the procedure is transparent but the absence of data on the shape distribution means that it cannot be tuned for selection of subgroups as in this study.

### Image orientation

While the outline of an object is a primary guide to its identification (Spröte et al., 2016; Elder, 2018) certain orientations are especially informative. For trochoidally-coiled taxa like *Truncorotalia* a preferred position is in the plane of the coiling axis: this encapsulates much of the coiling geometry including rate of whorl translation (height of early whorls), gross radial/axial dimensions and the axial extension of late-formed chambers. Spiral orientation is one normal to the coiling axis which provides a sectional or planar view at the latest stage of growth. While the outline is less informative than in axial orientation the interior view is valuable as it shows the full growth sequence of chambers and their shape. Umbilical orientation is a 180° rotation of the specimen from the spiral; as its outline is a reversed copy of that in spiral orientation it is redundant in this shape analysis. Its interior view is restricted to chambers in the outer whorl; the shape of the aperture is often obscured. Imagery of shells in this orientation was preferred by Elder et al. (2018) and Hsiang et al. (2019). Of the 287 images in the latter study of *Truncorotalia crassaformis* at http://endlessforams.org/ 261 are in that position.; Al-Sabouni et al. (2018, p. 521) noted in an AI-directed study that “The taxonomically more informative umbilical side was typically oriented upwards…”). This study suggests that if a single orientation is to be analysed for exemplars the axial is preferable. Some of the inconsistencies in identification reported byAl-Sabouni et al. (2018) might be due to the use of umbilical views.

## Conclusions

Planktonic foraminiferal taxonomy, like others, is based on visual judgement of specimens yet the question of how to create representative imagery that facilitates the identification of taxa has been neglected. Possibly this is justified: the current taxonomy of c. 50 species, although curiously small relative to some zooplankton, is validated by the valuable proxy environmental data (Schiedel et al., 2018) it creates. Nonetheless, exemplar images are the workhorses of taxonomy: “images are more vivid and indelible than words” (Daston & Galison, 1992). Although their selection remains largely at the discretion of the taxonomist it is advocated that identification of morphotaxa will be enhanced by the adoption of methods less dependent on visual judgement. Here the problem is viewed as one of making training sets (exemplars) that capture important aspects of population variation, given that taxa have been created by visual judgement. Essentially the problem is to find methods which raise transparency in the selection of exemplars from morphospecies that have been recognized visually. The morphometric approach trialled here does not eliminate discretionary judgement but it is reduced relative to visual consensus judgements about exemplars by expert panels. The algorithms used here are a few of many solutions to a significant problem in taxonomy.

Shape is a powerful summary attribute of shell architecture that is easily quantified and applied to the selection of exemplars. Recognition of specimens near the center of shape variation for use as exemplars (C-exemplars) is promoted because their abundance suggests that they are likely to represent typical, best adapted architectures. As well as drawing attention to the magnitude of variation, presentation of outliers in the shape distribution (B-specimens) assists with identification of the ‘difficult specimens’ (Sharma et al. 2019) that particularly perplex novices. Of the standard orientations for trochoidally-coiled shells, the axial position, as judged from outline data, may be the most informative. Expectedly, future advances in parsing structures within the outline will further contribute to the selection of exemplars.

